# Soil Water Carrying Capacity for Vegetation

**DOI:** 10.1101/827576

**Authors:** Zhong-sheng Guo

## Abstract

Non-native plantation can effectively conserve soil and water and improve ecological environment,but soil desiccation occurs and then changes to severe desiccation in most forest and grass lands of water-limited regions, which causes soil degradation, vegetation decline and crop failure, so it is necessary to regulate the changed relationship between plant growth and soil water. However, there is a lack of a universally accepted theory to provide guidance of sustainable use of soil water resources and sustainable management of forest and sustainable produce of crop in such regions. The theory of regulating the relationship between plant growth and soil water in water-limited regions includes the Soil Water Resources Use Limit by Plant (SWRULP) and Soil Water Carrying Capacity for Vegetation (SWCCV). The SWCCV is the population in a plant population or density of indicator plants in a plant community when the soil water supply is equal to soil water consumption in the root zone and a given time, and changes with plant community type, location and time. The degree of coverage, productivity and benefits of a plant community when population quantity of an indicator plant equals SWCCV should be the theoretical basis for ensuring sustainable use of soil water resources and sustainable development of forest.

## 1. Introduction

In most parts of the world, human activities, such as overgrazing, deforestation, denudation and reclamation have greatly altered the type of vegetation that dominates the landscape. These have accompanied the demand for food, timber and biofuels due to local population increases, which historically have frequently occurred in water-limited regions such as the Loess Plateau of China. Intense and poorly managed agricultural practices have often caused a decline in the density of natural plant populations (Metcalfe and Kunin 2006). Consequently, the original vegetation has disappeared and there has been a decrease in the level of forest cover and in the ability of forests to maintain a balanced ecosystem. Such changes have led to severe soil and water loss and continual degradation of the natural environment on the Loess Plateau, which severely affects the health and security of forest and vegetation ecosystems and humans.

In order to conserve soil and water and improve the ecological environment and restore harmonious human–human, human–nature and human–society relationships (i.e. ecological civilization construction), since 1950, large-scale afforestation has been carried out on the Loess Plateau. This has been especially supported since 1978 by implementation of projects of the ‘Three-north Protection Forest’ that involves planting trees and establishing protection forest in north-eastern, north-western and northern regions of China to control soil and water loss, prevent wind erosion and fix sand and improve the ecological environment in which people live and work. The ‘Converting Farmland to Forest’ has also been implemented since 2000 and strengthened vegetation restoration. Consequently, forest area and coverage has dramatically increased, as has the efficiency of forest and vegetation in conserving soil and water. For example, the sediment discharge on the Loess Plateau was reduced from 1.6 billion tonnes per year in the 1970s to 0.3 billion tonnes per year in recent years, runoff has halved and the environment has improved. The efficiency of forest and vegetation accounts for more than half of the total efficiency.

In the process of vegetation restoration, tree species, selected for their capacity to extend deep roots and for fast growth, were planted at high initial planting densities to rapidly establish high degrees of ground cover and higher biomass and yields, and thereby to quickly realize ecological, economic and social benefits during vegetation restoration. It is advantageous that the roots of these plants grow quickly and thus they take up water from considerable soil depths, such as 5.0 m for 16-year-old Caragana (*Caragana korshinskii* Kom.) of the semiarid loess hilly region in Guyuan County, Ningxia Hui Autonomous Region of China and 22.4 m for 23-year-old Caragana (*C. microphylla* Lam.), another related species in Suide County, Shaanxi Province of China (Wang *et al* 2009). However, soil water mainly comes from precipitation; and the maximum infiltration depth (MID) and soil water supplies are limited in this region (Guo and Shao 2013). Thus, root depth can exceed the depth of soil water recharge from rainwater, leading to severe desiccation of soil in rooting soil layers (Wang *et al* 2009;Guo and Shao 2003). Consequently, the combination of increased water use by plants and low water recharge rates has led to soil deterioration, receding vegetation and crop failures on the Loess Plateau or other water-limited regions in the perennial artificial grass and forest land (Yang 1996;Li 2001;Chen et al 2007). Such soil deterioration can adversely affect ecosystem function and services and the stability of man-made forest and vegetation ecosystems, and consequently reduces the ecological, economic and societal benefits of forest and other plant communities. In turn, this suggests that the relationship between plant growth and soil water (RBPGSW) in these perennial artificial grass and forest lands is not balanced and should be controlled.

Managing an ecological system to maximize the benefits of vegetation requires establishing a balance between soil water supply and soil water consumption over a long time in water-limited regions. This is because a rapid increase in density and the degree of coverage of forest and/or other plant communities not only effectively increases coverage and reduces runoff and erosion (Bosch and Hewlett 1982;Khanna et al 1999;Gardiol *et al* 2003;Dijk and Bruijnzeel 2003;Anderegg *et al* 2012) but also reduces soil water supply and increases soil water consumption by vegetation and evapotranspiration (Kyushik *et al* 2005), which can lead to imbalance in the RBPGSW. Therefore, there should be limits to vegetation restoration in a region where the natural resources (including water, soil water, soil nutrients or land resources) are scarce. The limit would depend on the capacity of the available natural resources in an ecosystem to support vegetation, i.e., land carrying capacity for vegetation or vegetation carrying capacity (Guo and Shao 2003).

Water is the main factor affecting vegetation restoration in most water-limited regions. The carrying capacity of land for vegetation in this region is soil water carrying capacity for vegetation (SWCCV)(Guo *et al* 2002;Guo and Shao 2003) – the ability of soil water resources to support vegetation. Therefore, the balance between the consumption and supply of water to the soil should be considered when restoring vegetation cover in order to realize the goal of soil and water conservation and sustainable use of soil water resources.

The Loess Plateau has the most serious soil and water losses in the world. It is located in the centre of China, and has an area of 642 000 km^2^ of which 454 000 km^2^ experiences soil and water loss and has scarce water resources. Soil in this region is very deep, in the range of 30–80 m from the surface (Zhu *et al* 1983), and the groundwater table is also deep (Yang and Shao 2000). Without irrigation, the best measure to solve issues of soil degradation and vegetation decline is to regulate the RBPGSW by reducing the population quantity of indicator plants in a plant community to match SWCCV on the Loess Plateau, thus balancing the soil water recharge and soil water consumption in plantations (Guo an shao 2003).

Drought is a recurring natural phenomenon. The complex nature and widespread impact of drought on forest and grass land with high coverage and production – driven by artificial vegetation consuming more than a permissible quantity of soil water resources in water-limited regions – means that regulating the RBPGSW is needed to maintain soil water consumption of restoring vegetation at levels that sustainably use the soil water resources. However, there is a lack of a universally accepted theory to provide guidance for regulating the RBPGSW in forest and grass land in these regions. We aimed to develop the theory for regulating the RBPGSW in forest and grass land in water-limited regions, including (1) the concepts of soil water resources, (2) soil water resource use limits for plants, (3) SWCCV, (4) basic laws of SWCCV and (5) the use of SWCCV in the sustainable management of forest and vegetation.

## 2. Soil water resources

Soil water is the water existing in the soil space. Soil water cannot be transferred by humans from one place to another but can be used by plants. Soil can hold much water and this is sometimes termed a soil reservoir. The amount of water held in the soil is the soil water resource. Soil water resources come (Budagovski 1985)after the overall soil moistening (Lvovich,1980). Soil water resources is the water stored in the soil from surface soil to the groundwater table-generalized soil water resources or the water stored in the root zone soil-Soil water resources in narrow sense, and the dynamic soil water resources - the antecedent soil storage in the root zone plus the soil water supply from precipitation in the plant growing period or a year for evergreen plants – because soil water from precipitation in the growing season can be taken up immediately by live plants and influence their growth.

Plant evapotranspiration including plant transpiration and film water evaporation on the surfaces of leaves, branches and stems when raining and after a rain event is a hydrological effect that induces soil suction. With water absorption by roots and evapotranspiration, soil water content and soil water potential are reduced but soil water suction is increased. When soil water content in every soil layer is reduced to the wilting coefficient, the suction of soil particles to water exceeds that of plants to soil water and the water stored in all soil layers cannot be further taken up by plants (i.e. it is plant-unavailable). When soil water content in every soil layer is higher than wilting coefficient, the suction of soil particle to water is lower than that of plants to soil water and so this part of soil water resources is plant-available and can be sucked by plant.

## 3. Soil Water Resource Use Limit by Plants

A tree or plant is a complex organism with a series of regulatory mechanisms to keep vital systems operating within appropriate restrictions, and with mechanisms to repair damage that may occur when these limits are exceeded. A tree or plant transports water from the soil to the atmosphere along a water potential gradient; its survival depends on maintenance of this transport system (Anderegg *et al* 2012). Plants also have some self-regulation function in the opening degree of stomata in leaves and the number of leaves retained during water stress; however, this self-adjustment is limited and cannot meet the need of regulating RBPGSW in the process of plant growth in some extreme conditions, such as severe drought and hot days on the Loess Plateau. While a plant grows, individual size expands, Canopies effect on soil water supply and soil water consumption strengthens and roots deepen but soil water resources in artificial forest and grass land often decline in water-limited regions, even if there are some increases after rain events, and then root water uptake declines. Water stress begins when transpiration demand exceeds root water uptake, resulting in a loss of turgor. Subsequent short- and long-term responses include declines in cell enlargement and leaf expansion rate, reduced photosynthesis and transpiration, and alterations in phenology, senescence, carbon allocation and ultimately reduced yield and water use efficiency. When soil water resources reduce to the degree, that results in severe soil drought, and finally leads to soil degradation, plant growth ceases and vegetation recedes or even dies in artificial forest and grass land.

The RBPGSW is important for the prevention of further soil drying and soil degradation and the sustainable use of soil water resources and sustainable management of forest vegetation in water-limited regions. In practice, regulating the RBPGSW is not required once soil drought happens in forest and grass land.

There are two important soil water deficit criteria: Soil Water Resources Use Limit by Plants (SWRULP)(Guo 2010,2014). SWRULP can be defined as the soil water storage in the MID when soil water content in all soil layers within the MID equals the wilting coefficient and roots can no longer withdraw water within the MID.

Infiltration depth is one of the most important indexes for estimating soil water deficit criteria. The infiltration depth for one rain event equals the distance from the surface to the crossover point between the two respective soil water distribution curves of soil water with soil depth before and after the rain event (Fig. 1a). The MID will occur after a continuous heavy rainfall event and a long-term cumulative infiltration process, and can be determined by a series of two-curve methods (Fig. 1b). Such as in the Shanghuang Ecological Experimental Station, Annual precipitation ranged from 284.3 mm in 1986 to 634.7 mm in 1984. For example, the MID was 290 cm, which happened on 1 August 2004 after some heavy rain in August 2003– 55.0 mm on 1 August, 45.7 mm on 25 August and 56.4 mm on 26 August for Caragana shrub land in the semiarid loess hilly region of the Loess Plateau.

**Fig. 1.**
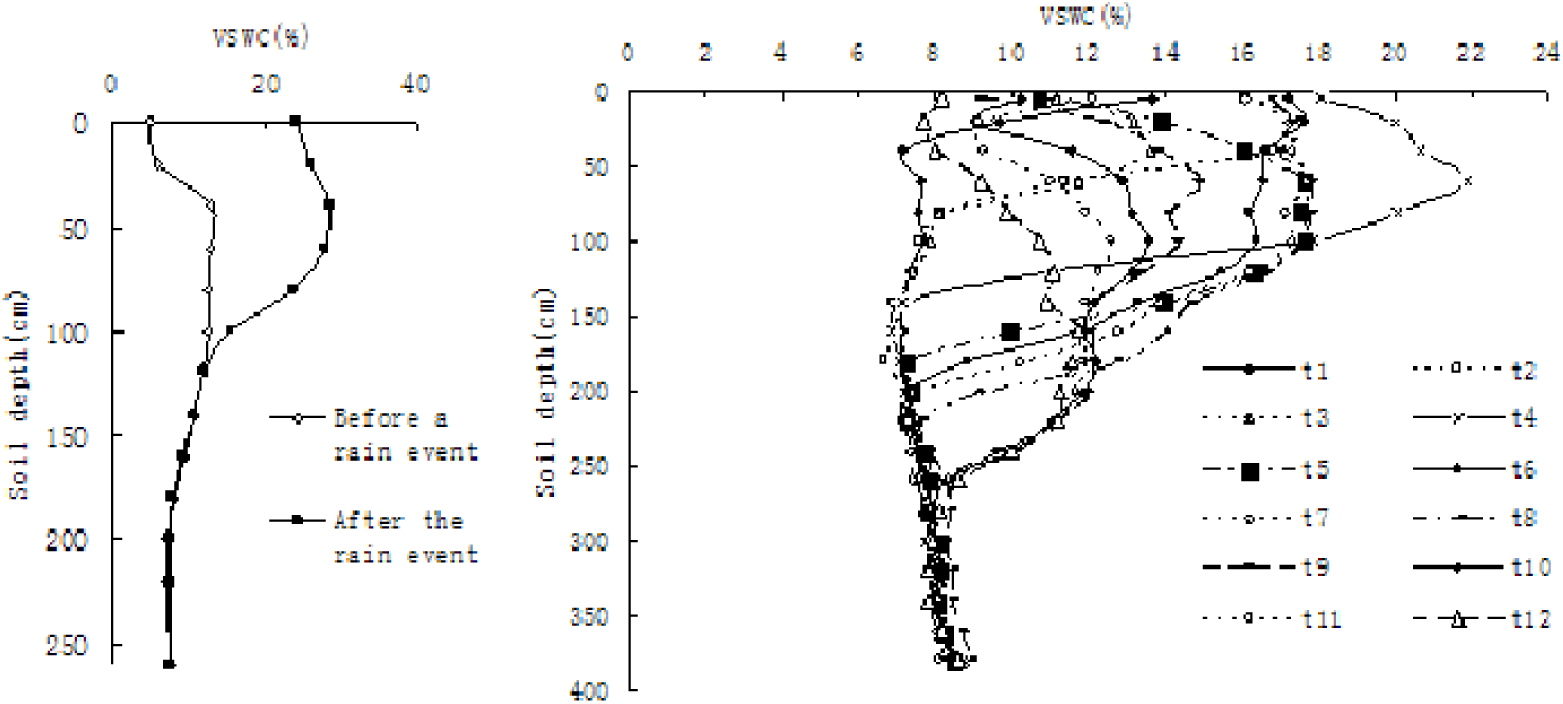
Two-curve method for estimating infiltration depth. a, infiltration depth and soil water supply for one rain event. b, The maximal infiltration depth is estimated by a series of two-curve methods in the soil under Caragana shrubland in the semiarid loess hilly region of China. VSWC is Volumetric soil water content.

The root vertical distribution is another important index for estimating soil water deficit criteria. For example, the majority of Caragana root biomass was distributed in the 0–200 cm soil layer even though roots extended to 5.0 m, MID was 2.9 m, SWRULP was 222.8 mm and LSWR was 405.7 mm in 16-year-old Caragana shrub land of the semiarid loess hilly region (Fig. 2). The amount of water carried from the soil through plants to the atmosphere depends on weather, plant growth and soil water conditions. If the soil water supply is smaller than soil water consumption then soil water resources will reduce and eventually reach the LSWR, then plants will die.

**Fig. 2.**
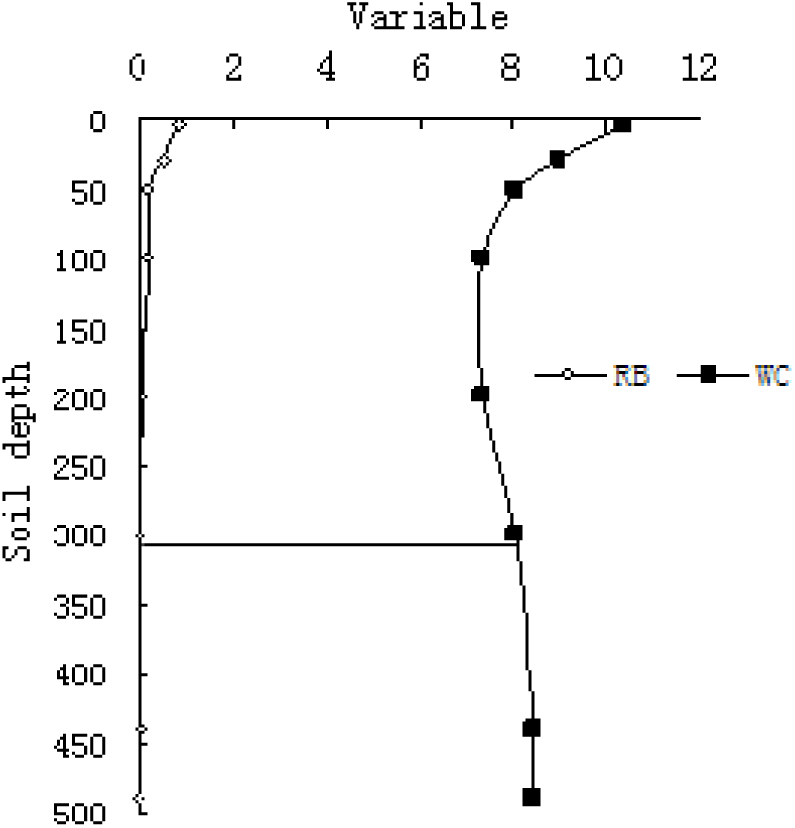
The relationships among root distribution depth (RDD), wilting coefficient (WC), MID, SWRULP and LSWR in 16-year-old Caragana shrubland in the semiarid loess hilly region of China.

When the soil water resources in the MID equal the SWRULP, even if plants withdraw some water in soil layers deeper than the MID, the amount of root water uptake is small and will not meet the needs of plant transpiration. Therefore, this is the appropriate start time to regulate the RBPGSM. For example, the start time is the fifth year after caragana sowing in the degradation and receding vegetation. The amount of regulation is the difference between plant density and SWCCV, because SWCCV is the measurement of sustainable use of soil water resources by plants. When existing plant density exceeds the SWCCV and the difference between soil water supply and soil water consumption is greater than zero then soil water deficit occurs and is exacerbated with time because soil water supply is reduced (Fig. 3a) and soil water consumption increased (Fig. 3b) with increasing density in a given period. This suggests that available soil water resources will not support the existing plant population and the population should be reduced. The difference between existing plant density and SWCCV indicates the amount of regulation required.

**Fig. 3.**
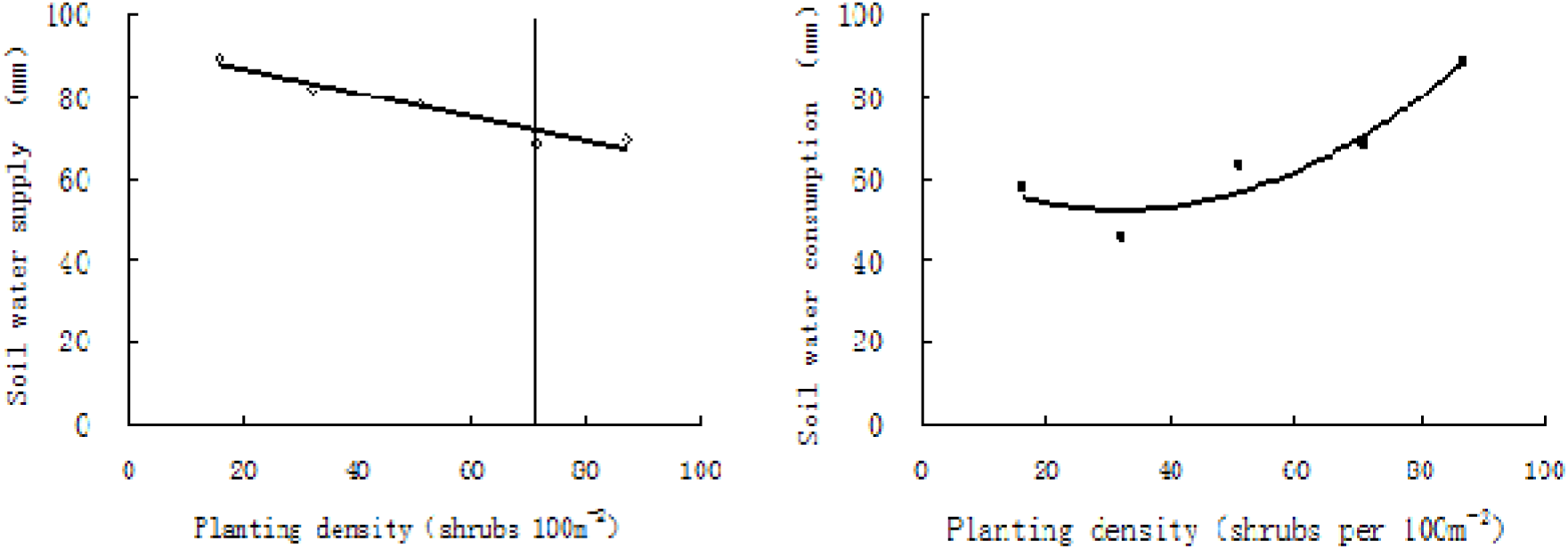
Changes in soil water supply and soil water consumption with planting density and SWCCV in Caragana shrubland of the semiarid loess hilly region, China. a, The change in soil water supply with planting density and SWCCV; b, The change in Soil water consumption with planting density and SWCCV.

## 4. Soil Water Carrying Capacity for Vegetation

The idea of carrying capacity has its origin in the doctrine of Malthus (Steiguer 1995). The term carrying capacity was first used by range managers (Price 1999) and U.S. Department of Agriculture researchers (Young 1998). After Raymond Pearl and Lowell J. Reed proposed logistic equations in 1920, Odum (1953) equated the term carrying capacity with the constant K in logistic equations (Price 1999;Young 1998), see equation 1:

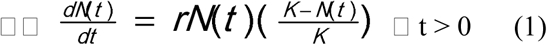

Where N(t) is density at time t, the population per unit area at time t, r is the intrinsic growth rate, r > 0 and K is an asymptote (the carrying capacity) with K > 0.

### The concept of SWCCV

Since the 1960s, soil drought has occurred in most of the perennial grass, forest and crop land in the Loess Plateau in China. The concepts of vegetation carrying capacity, the ability of land resources to support vegetation appeared (Guo *et al*, 2002). Vegetation carrying capacity includes space vegetation carrying capacity, the carrying capacity of land or space resources for vegetation, soil water carrying capacity for vegetation (SWCCV) and soil nutrient carrying capacity for vegetation (SNCCV). SNCCV can be subdivided into soil nitrogen or potassium or phosphorus carrying capacity for vegetation and so on.

Vegetation includes different plant populations or communities. In nature, no plant community is formed by a single plant species and form a plant populations, instead many plant species live together and form the community to use space, soil water or soil nutrient. Although any one species in a plant community can express SWCCV in this situation in theory, but the different plant species differ in their positions and roles in the community. Constructive species for natural vegetation and principal or purpose species of trees or grasses are selected drought resistant plant species – principal species are the main afforestation tree species and account for the majority of the trees planted in a region, and purpose species of trees or grasses are those cultured or managed by people in a region. Generally, principal species of trees or grasses are also purpose species in a region; for example, Caragana in the Loess Plateau is the regulated tree species.

Planting trees is a purposeful human activity to meet the need of different people; Caragana is planted where vegetation is sparse and soil erosion is serious. Such plant species are often exotic; sensitive to water deficits; play the most important roles in preventing wind erosion, fixing sand and controlling soil,and population quantity of Caragana play the most important roles in preventing wind erosion, fixing sand and controlling soil and water loss in plant communities; and become the goal of cultivation, management and regulation; and become the goal of cultivation, management and regulation.. Thus, an indicator (plant) species is the representative of a plant community to express SWCCV – either a constructive species for natural vegetation, or a principal or purpose species for artificial vegetation. The SWCCV can be defined as the maximum plant population quantity (absolute index) or plant density (relative index) of an indicator species in a plant community when soil water consumption is equal to soil water supply in the root-zone soil layers. The value of SWCCV is a function of indicator species, connected with plant community type, time and sites. The value of SWCCV is a function of indicator species, connected with plant community type, time and sites (Guo 2014).

### 4.1 Choice of scale

The choice of scale is important for calculating SWCCV. In fact, different time-scales can be used to estimate SWCCV. SWCCV is meaningless on such short time-scales as 24 hours or a day, and two years or more is too long. Minimum death days or minimum cripple days is an appropriate time-scale to estimate soil water supply, soil water consumption and then determine SWCCV, and then judge whether population quantity or plant density exceeds SWCCV. Minimum lethal days is the most suitable time scale.

Different space-scales can also be used to estimate soil water supply, such as 1 or 10 km^2^. However, a crustal block or slope is a suitable space scale to compute SWCCV because there are changes in such environmental factors as radiation, precipitation, wind and temperature with increasing space scale, which may change vegetation type. If the space scale is too large, the environmental factors and the community type will change and so influence SWCCV.

### 4.2 Methods of evaluating SWCCV

There are equations to estimate SWCCV, such as classic carrying capacity models (Cohen 1995), general models of population growth (Price 1999) and physically-based process models (Xia and Shao 2008)and soil water–plant density models; however, soil water–plant density models are best.

#### 4.2.1 Classic SWCCV model

Classic carrying capacity models were proposed by Albrecht Penck. In 1925, he stated a simple formula that has been widely used (Cohen 1995). According to classic carrying capacity models, the classic SWCCV model is presented in the following form:

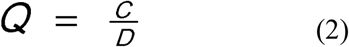

Where, C is soil water resources per unit area and D is individual water requirement. Ma *et al*. estimated SWCCV for eight tree species using this formula in a dry and warm valley of Yunnan, China (Ma *et al* 2001). The minimum water consumption that an individual plant consumes water over a season or year when growing in a normal and healthy way should be used as the individual water requirement – because water consumption of an individual plant over a year changes with weather, plant growth and soil water conditions. If the minimum water consumption is used as the individual plant water requirement, which is suitable for saving water and precise soil water management in forest and grass land even the individual water requirement index, that is, the minimum water consumption is too high.

#### 4.2.2 General population growth model

This model was proposed in 1920 by Pearl and Reed, who reasoned that there must be an absolute limit beyond which further population growth would be impossible:

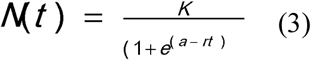

Where N_(t)_, K and r are the same as equation 1; and parameters a and e are constants. Given a population at different times, and at least three sets of populations and relative time data, then the carrying capacity can be obtained using equation 3. Because the equation has no soil water resources term,it is difficult to obtain the SWCCV.

#### 4.2.3 Soil water–plant density model

If we established a set of different population quantities (or densities) of indicator plants, E_1_, E_2_, E_3_…, E_n_ at the same condition and the same plot areas (e.g. 5 m × 20 m plots for Caragana in the Loess Plateau), so these plots have the same radiation, temperature, annual precipitation and its distribution in a season or a year, slope, slope aspect, slope position and soil type would be almost the same. However, they would have differing population quantity or density and so different effects on soil water supply and soil water consumption. The precipitation, throughfall, runoff, plant growth, deep seepage and soil water changes with soil depth and time in the root zone are measurable variables, thus the soil water supply and soil water consumption for the different population quantities or densities on a growing season or an annual basis can be measured, collected and statistically analysed. Soil water supply reduces with increased planting density (Fig. 3a); and the relationship between soil water consumption and population quantity or density can be described by a parabolic equation (Fig. 3b). The quantitative relationship between soil water supply or soil water consumption and population quantity can be established using a least squares method. Generally, the soil water and plant density model can be expressed in the following form (Fig. 2):

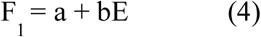

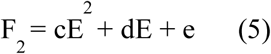

Where F_1_ is soil water supply, E is population quantity (or plant density), F_2_ is soil water consumption and parameters a–e are constants (Guo and Shao 2004). SWCCV can be determined by combining equations 4 and 5, with the positive solution being SWCCV. During 2002 to the present, we studied Caragana shrub land planted by sowing in fish-scale pits on slopes in the semiarid Loess Plateau. For example in 2002 in 16-year-old Caragana shrub land (Guo 2014), F_1_ = 92.494 – 0.2913E (see Fig. 3a), F_2_ = 0.0118E^2^ – 0.7575E + 64.759 (see Fig. 3b) and SWCCV is 72 bushes per 100 m^2^ or 7200 bushes per ha. Stem number, S, changes with plant density E, and the S and E relationship is: S = 42.88E – 62783, with R^2^ = 0.9632.

#### 4.2.4 Physically-based process model

Xia and Shao proposed a process model (Xia and Shao 2008). They determined SWCCV according to the number of days that soil water content equaled the wilting point was less than the maximum number of days. This equation has many parameters and calibrating every parameter leads to error. Additionally, sclerophyllous plants such as Caragana adapt to severe dry conditions through change of leaf colour and early leaf fall. The SWCCV estimated by this method does not ensure that the soil water supply equals soil water consumption in a growing season or a year when plant density equals SWCCV, and thus this model needs to be improved.

Currently, the soil water and plant density model is the best model to estimate SWCCV. Once a set of different density experimental plots is established, SWCCV can be estimated at different tree ages and annual precipitation. The parameters a–e change with time and site.

### 4.3 The basic law of SWCCV

The value of SWCCV changes with year because precipitation changed with year, such as the precipitation in Shanghuang Ecoexperimental Station in the semiarid region (Ning xia Guyuan of China) varies from 634.7 mm in 1984 to 259.9 mm, which influenced the soil water supply. At the same time when trees grow, and the forest canopy, annual precipitation, soil water supply and soil water consumption at different population quantities or densities change on an annual basis. In 2002, the relationships between soil water supply and E, and soil water consumption and E, are shown in Fig. 3a and b, respectively. Combining and solving the simultaneous equations, the value of SWCCV for 16-year-old Caragana was 7200 bushs per ha because Caragana was clustered in fish-scale pits. The annual precipitation of 623.3 mm in 2003 was close to the record maximum, and was considered as a 19-year event. Caragana is a deciduous species and soil water storage at the end of the growing season exceeded that at the beginning for a plant density of 8700 bushes per ha (i.e. the maximal density of the year-old experimental Caragana shrub land), thus the soil water supply exceeded soil water consumption in the root zone at that time. Because there were no Caragana deaths in the plot, this suggested that the SWCCV was 8700 bushes per ha in this case. The value of SWCCV for 18-year-old Caragana was less than 1600 bushes per ha (because this was the minimal experimental density for this shrub land) in 2004 because the annual precipitation of 328.3 mm in 2004 was low, and soil water supply was less than soil water consumption in the root zone and there was no recorded Caragana death in the plot. In Caragana shrub land planted in 2002 by broadcast sowing, the SWCCV for 10-year-old Caragana was 480 000 stems per ha in 2011; in 2012, this was 420 000 stems per ha for 11-year-old Caragana; and, in 2013, 420 000 stems per ha for 12-year-old Caragana. Thus, the RBSWPG for 10–12-year-old Caragana did not require regulation because the soil water resources in the year could support the maximum experimental density, suggesting that value of the SWCCV changed with time (year) at the same site (Guo 2014).

The value of SWCCV varies with indicator species or vegetation types at the same place during the same period. This point is supported by the study results of eight indicator trees species using the classic carrying capacity calculation formula (equation 2) in a dry and warm valley of Yunnan, China. The value of SWCCV differed across these species (Ma *et al* 2001).

The value of SWCCV changes with location or site because soil water resources mainly come from and are closely related to precipitation (Koster *et al* 2004;Wang *et al* 2006), and the annual precipitation, suitable plant species and the requirements of economic and social development for forest and vegetation restoration differ with location. The soil water supply and evapotranspiration in forest land differ with vegetation type within the same period, so the soil water resources vary with location in water-limited regions. This is supported by four reports from China: in the Inner Mongolia Autonomous Region (Tian *et al* 2008), in Shenmu county of Shanxi (Xia and Shao,2008), in the Donggou Valley of Mizi (Wang and Shao 2012) and in Diedie Gully Valley in the Ningxia Hui Autonomous Region (Liu *et al* 2009, jia et al,2019).

### 4.4 Use of SWCCV in practice

#### 4.4.1. Fundamentals required for determining the amount of trees to be cut

The soil water resources in Caragana shrub land are reduced with increased planting density and time (forest age) under rain-fed conditions, except during wet years. When the density of the indicator species, Caragana, exceeds its SWCCV value, soil water content under the shrub land is reduced and threatens the health and stability of the shrub vegetation ecosystem. When the soil water resources drop to the SWRULP, soil water seriously impedes plant growth (Guo 2010b). Thus, the RBPGSW needs to be regulated using SWCCV in water-limited regions. The time to start regulating the RBPGSW is at the fifth year for Caragana in the semiarid Loess Plateau (Guo and Li 2009b).

If the population quantity or density of an indicator or purpose species of tree in a plantation exceeds the SWCCV value, the higher ecological, economic and social benefits and productivity in the plant community will be obtained at the expense of the environment (more serious soil drying, soil degradation and vegetation decline). Even if a soil water deficit in the plant community or forest vegetation does not immediately destroy the ecosystem, this situation will not well sustain the vegetation. If density of the indicator species is less than the SWCCV value, the plant community does not make the most of natural resources. Thus, the present productivity, ecological, economic and social benefits of the plant community does not reflect the maximum services and benefits the rain-fed ecosystem function can provide and wastes the soil water resources.

SWCCV is the foundation to determine the amount of regulation needed. The cover degree is an important index to express the function of vegetation in reducing raindrop dynamic energy and wind velocity near the ground and conserving soil and water. The degree of cover (Fig. 4a) and productivity (Fig. 4b) in Caragana shrub land increases with planting density. Because the main erosive force is runoff in the Loess Plateau, when Caragana density exceeds the SWCCV value there is higher cover degree (Fig. 4a) and canopy interception (Fig. 5a), as well as lower runoff (Fig. 5b) and then lower soil loss (Fig. 5c). However, this is at the expense of the soil water environment because soil water consumption exceeds water supply from rainfall when planting density is more than SWCCV, which is not good for sustainable use of soil water resources and sustainable management of forest land. Keeping the planting density of Caragana at the level of SWCCV is required to balance soil water supply from rain and the plants’ water requirements and make the most of soil water resources. In many cases, sustainable management of forest must be applied when the density of the indicator species in a plant community equals SWCCV in order to avoid soil degradation and vegetation recession. The amount of trees that should be cut when regulating equals the existing density minus the SWCCV.

**Fig. 4.**
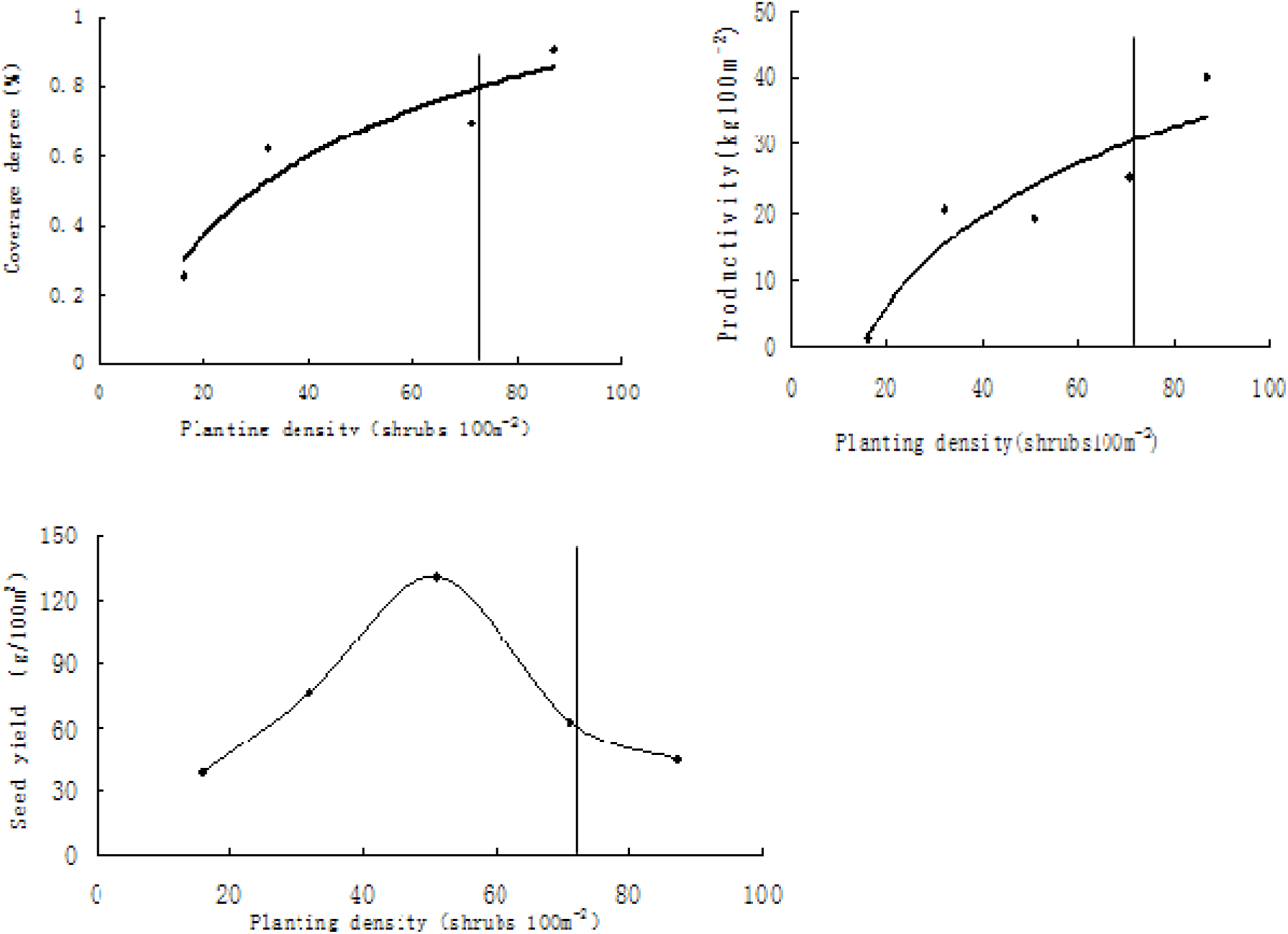
Changes in coverage degree, pruductivity and seed yield with planting density and SWCCV in Caragana shrubland of the semiarid loess hilly region, China. a, The change in coverage degree with planting density and SWCCV. b, The change in Pruductivity with planting density and SWCCV. c, The change in seed yield with planting density and SWCCV.

**Fig. 5.**
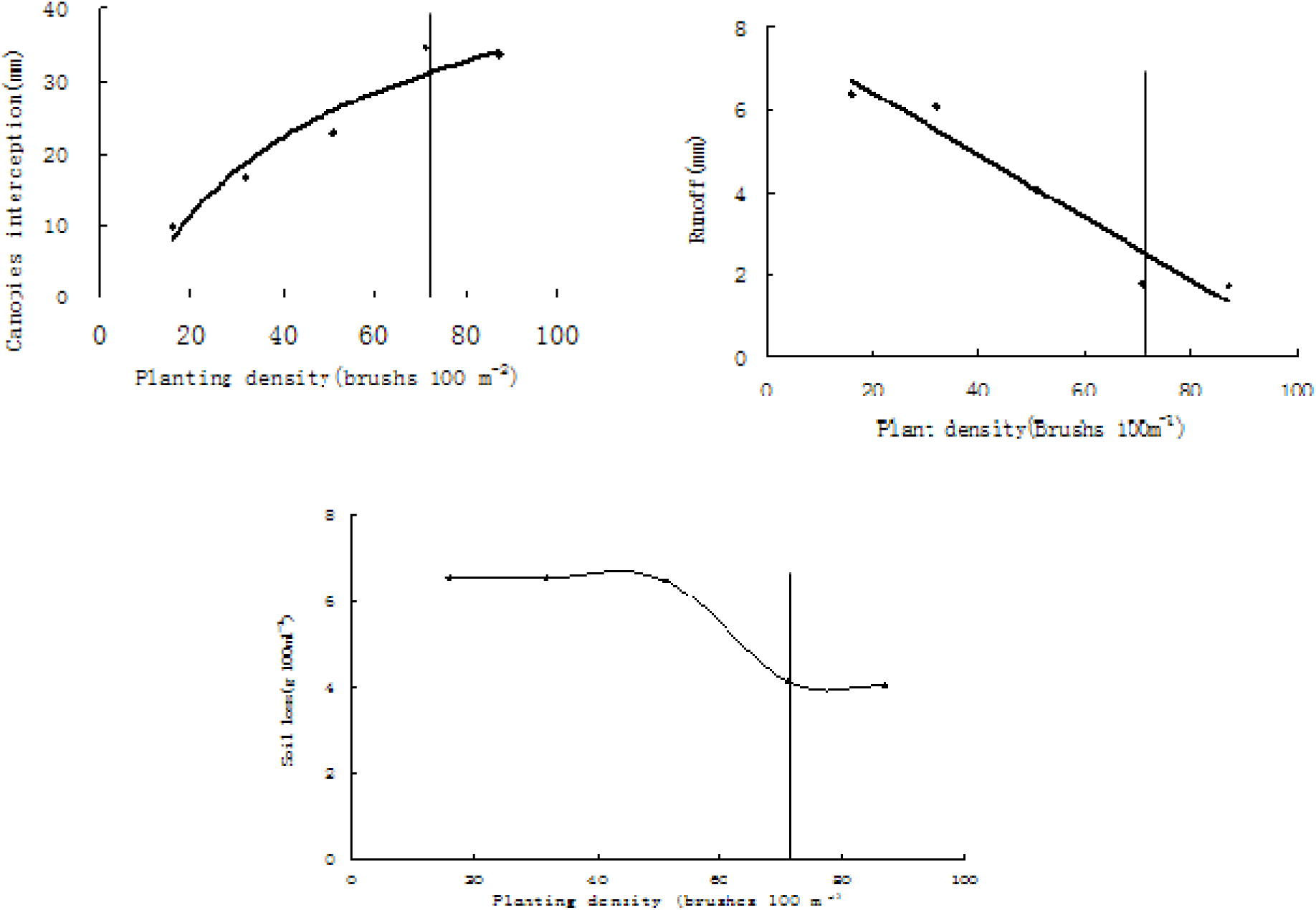
Changes in canopy interception, runoff, soil loss with planting density and SWCCV in the Caragana shrubland in the semiarid loess hilly region of China. a, The change in canopy interception with planting density and SWCCV; b, The change in runoff with planting density and SWCCV; c, The change in soil loss with planting density and SWCCV.

#### 4.4.2 Theoretical basis to determine criteria and indicators for sustainable forest management

Since the beginning of the 1990s, an enhanced understanding of sustainable forest management (SFM) has entered forest policy worldwide, with the concept of criteria and indicators (C&I) as one cornerstone for implementation (Wolfslehner *et al* 2005). Over the previous decade, SFM has become a highly relevant topic in both forest and environmental policy (Sheppard and Meitner 2005). Although much effort in SFM has focused on defining C&I for measuring sustainability (Wolfslehner and Vacik 2011), there is no universally acknowledged method for its determination (Wolfslehner *et al* 2005;Sheppard and Meitner 2005;Wolfslehner and Vacik 2011).

SWCCV is the foundation to determine the C & I for SFM. When the density of indicator species in a plant community equals the SWCCV value, the plant community makes the most of the soil water resources, and conditions of the community such as the appearance, crown coverage, productivity, constituents and carbon-fixing capacity should be indexes or the theoretical basis to determine C & I for sustainable management of the vegetation. The cover degree, biodiversity, productivity, biomass and its components and carbon-fixing capacity when the density of indicator species in the plant community equals the SWCCV value and the requirement of stakeholders should be combined to determine C & I for SFM.

This will enable maintaining and enhancing forest resources and their contribution to global carbon cycles, forest ecosystem health and vitality, productive functions of forests (wood and non-wood) and biological diversity in forest ecosystems because SFM depends not only on natural resources’ carrying capacity but also the support and input of a wide range of stakeholders (Sheppard and Meitner 2005;Wolfslehner and Vacik 2011; Masiero *et al*). For example, in the Loess Plateau of China, the value of SWCCV for 16-year-old Caragana is 7200 bushes per ha. When Caragana density is equal to The carrying capacity (i.e. 7200 bushes per ha), the cover degree (the ratio of the total area of Caragana canopies to experimental plot area or land area) of the 16-year-old Caragana is 80% (Fig. 4a), which should be the suitable vegetation restoration limit.

At this point, the biomass production is 1400 kg (dry weight) per ha per year (Fig. 4b), which should be the appropriate productivity, and the carbon content in the 1400 kg of biomass is the carbon-fixation capacity of Caragana. The seed yield of forest is smaller when the planting density equals SWCCV (Fig. 4c). Thus, SWCCV is the foundation to determine the C&I for sustainable Caragana shrub land management in the semiarid Loess Plateau.

## 5. Conclusion

Rapid global economic development and increase in population in the past century has led to unprecedented consumption of natural resources to satisfy the growing demands for food, fibre, energy and water in water-limited regions. These natural assets were perceived as free and limitless for many decades. Nowadays, with scientific, technological and economic development, increasing public awareness of the value of soil, water and forest resources and the growing need for environment quality has gradually led governments to adapt their policies and strategies to match sustainability goals. This has meant dealing with the overuse of soil water resources, soil degradation and vegetation decline in the process of soil and water conservation and vegetation restoration in water-limited regions. Sustainable use of natural resources and keeping the environment sustainable in water-limited regions is the basis for sustainable development of social economy and human consensus because the area of water-limited regions accounts for the most of global land and supports a numerous and ever growing population.

Carrying capacity is the measure of sustainable use of natural resources. SWCCV is the core issue for forest and vegetation restoration, sustainable use of soil water resources, SFM and restoration of a harmonious relationship between humans and nature in water-limited regions such as the Loess Plateau. We should plant trees and sustainable management forests in soil and water loss regions to conserve soil and water and improve the environment. In water-limited regions, soil water resources are reduced with growth of forest even with a sudden increases of soil water resources after rain events. When the soil water resources equal the SWRULP, soil water severely constrains plant growth. When this occurs, SWCCV should be estimated – on which the RBPGSW can be regulated.

SWCCV is the ability of soil water resources to support vegetation and can be defined as the maximum plant population quantity (absolute index) or plant density (relative index) of indicator (plant) species in a plant community when soil water consumption is equal to soil water supply in the root zone on an annual basis in a water-limited region. The value of the SWCCV is a function of plant community (indicated by indicator species), time (expressed in years) and location. When the density of indicator species in a plant community is more than the SWCCV value, the RBSWPG should be regulated. When the density of indicator species in a plant community equals the SWCCV value, the conditions of the community such as the appearance, crown coverage, productivity and its constituents and carbon-fixing capacity should be indexes or the theoretical basis to determine C&I for sustainable management of forest and vegetation and sustainable produce of crop. The SWRULP and SWCCV provide a scientific basis for sustainable use of soil water resources in the process of vegetation restoration and determining C&I for SFM in water-limited regions.

## Acknowledgement

This study was supported by National key Research & Development plan (Project No. 2016YFC0501702)and the National Science Fund of China (Project Nos 41071193 and 41271539), We thank International Science Editing (http://www.internationalscienceediting.com) for editing this manuscript.

